# Evaluation of data-driven MEG analysis approaches for the extraction of fMRI resting state networks

**DOI:** 10.1101/2021.04.29.441916

**Authors:** Esther Annegret Pelzer, Abhinav Sharma, Esther Florin

**Affiliations:** Institute of Clinical Neuroscience and Medical Psychology, Medical Faculty, Heinrich-Heine University Düsseldorf; Max-Planck-Institute for Metabolism Research Cologne, Germany, Gleueler Str. 50, 50931 Cologne, Germany

**Keywords:** MEG, fMRI, Envelope correlation, phase-amplitude coupling, ICA

## Abstract

The electrophysiological basis of resting state networks (RSN) is still under debate. In particular, no principled mechanism has been determined that is capable of explaining all RSN equally well. While magnetoencephalography (MEG) and electroencephalography (EEG) are the methods of choice to determine the electrophysiological basis of RSN, no standard analysis pipeline of RSN yet exists. In this paper, we compare the two main existing data-driven analysis strategies for extracting resting state networks from MEG data. The first approach extracts RSN through an independent component analysis (ICA) of the Hilbert envelope in different frequency bands. The second approach uses phase –amplitude coupling to determine the RSN. To evaluate the performance of these approaches, we compare the MEG-RSN to the functional magnetic resonance imaging (fMRI)-RSN from the same subjects.

Overall, it was possible to extract the canonical fMRI RSN with MEG. The approach based on phase-amplitude coupling yielded the best correspondence to the fMRI-RSN. The Hilbert envelope-ICA produced different dominant frequency-bands underlying RSN for different ICA runs, suggesting the absence of a single dominant frequency underlying the RSN. Our results also suggest that individual RSN are not characterized by one single dominant frequency. Instead, the resting state networks seem to be based on a combination of the delta/theta phase and gamma amplitude.

## 1. Introduction

During the past decade, a well-reproducible connectivity map of brain activity during rest has been identified and thoroughly investigated in healthy humans using functional magnetic resonance imaging (fMRI) (Damoiseaux *et al*., 2006). Resting state analysis has gained increasing popularity in neuroscience because the data are relatively easy to acquire and do not depend on a task. Using MEG, recent studies have started investigating the temporal dynamics of resting state networks (Baker *et al*., 2014; Vidaurre *et al*., 2016). However, despite these advances, the electrophysiological underpinnings of the *canonical* fMRI RSNs are still not completely understood.

In the present paper we aim at providing a rigorous comparison of different approaches to extract these canonical fMRI resting state networks from MEG data. We do so by comparing RSNs extracted from resting state recordings for the same subjects in the MEG and fMRI. Using RSNs from the same subjects allows us to eliminate potential biases stemming from individual variability in fMRI RSNs. Because the literature considers the networks extracted from fMRI as the gold-standard, we take those as the benchmark and evaluate the MEG approaches according to their ability to match the fMRI results in the same subjects. In our comparison, we restrict our attention to data-driven approaches. While seed-based approaches can also be found in the literature (Hillebrand *et al*., 2012; Hipp *et al*., 2012; Marzetti *et al*., 2013), the seed choice adds a degree of freedom into the analysis that is difficult to control for. The ultimate goal of our study is to provide researchers with guidelines on how to extract canonical resting state networks in a data-driven manner from MEG data.

The MEG-RSN literature has provided conflicting findings on the main frequencies underlying individual RSN. For example, seed-based envelope correlation ascribes the dominant frequency ranges for the default mode network (DMN) to theta and alpha and for the dorsal attention network (DAN) the alpha and beta ranges (de Pasquale *et al*., 2010). In contrast, a data-driven approach based on independent component analysis (ICA) of frequency-specific envelopes (Envelope-ICA approach, Brookes et al. (2011)) obtained the best correspondence for the DMN within the alpha band and for the DAN within the beta band. In the present paper we investigate the role of RSN extraction techniques for potentially explaining some of these differences.

The phase lag index has also been proposed to obtain MEG-RSN (Stam *et al*., 2007; Hillebrand *et al*., 2012; Marzetti *et al*., 2013). According to these studies, most of the functional connectivity in the tested RSNs is promoted through alpha and beta oscillations. Unfortunately, these phase-lag index studies have so far been limited to seed-based analysis and no purely data-driven approach exists for the purely phase-based resting state extractions. On the other hand, it was demonstrated with a data-driven approach that phase-amplitude coupling between a low-frequency phase and high-gamma amplitude can explain the formation of the fMRI resting state networks (Florin and Baillet, 2015) (megPAC approach) – thereby combining the information from amplitude and phase.

Within the present paper we compare the two different data-driven approaches to extract the canonical resting state networks from MEG data based on resting state recordings from the same subjects in the MEG and fMRI. Their advantage is that they can be applied without any a-priori assumptions on particular seed locations. Insights on the correspondence to fMRI resting state networks from such data-driven approaches will be generalizable for future MEG studies and therefore might provide important insights on the electrophysiological underpinnings of fMRI resting state networks.

## 2. Materials and Methods

We included 26 healthy right-handed, male subjects [age: 26.7+/-3.9 SD; Edinburgh Handedness index 88.6+/-20.7; mini mental status test: 29.8+/-0.5]. Three of these subjects had to be excluded due to movement artefacts or technical problems during data acquisition. Before data acquisition, all subjects gave their informed consent and were then included in the experiment in line with the ethical guidelines of the declaration of Helsinki (Ethics committee Cologne: 14-264, Ethics committee Düsseldorf: 5608R).

In the following, we will first explain the MEG data pre-processing and the resting state network extraction, before describing the fMRI resting state network extraction and our approach for comparing the two.

### 2.1 MEG data acquisition and pre-processing

The MEG resting state data were acquired in a 306 channel MEG (Elekta-Neuromag) system with a sampling rate of 2400 Hz and a 800 Hz anti-aliasing filter. In total, 30 minutes of resting state activity in the MEG was recorded in a seated position for each subject. Subjects were asked to rest with eyes open and to look onto a fixation cross to reduce eye movement. The fixation cross was printed on paper and placed in front of the subject. This analogue setup was used to exclude the possibility that the projector’s refresh rate would lead to further extraneous frequency components (Logothetis *et al*., 2009). The MEG data were acquired in blocks of 10 minutes so that subjects could move in between the 10-minute blocks. Each 10-minute block should be long enough to capture the basic resting state fluctuations as was recently recommended for MEG measurements (Liuzzi *et al*., 2017). To monitor the subject’s head position 4 head-positioning coils were taped to the subject’s scalp. The positions of the coils were measured relative to the subject’s head using a 3-D digitizer system (Polhemus Isotrack). For anatomical co-registration with MRI, about 100 additional scalp points on the subject’s scalp were also digitized. In addition to the MEG, we simultaneously recorded an electrocardiogram (ECG) and electrooculogram (EOG).

After data acquisition, the preprocessing of the MEG data was done with standard processes implemented in brainstorm ((Tadel *et al*., 2011), https://neuroimage.usc.edu/brainstorm/). After the recording the line noise and its harmonics were removed (50, 100, 150, 200, 250 Hz) and sensors with high noise levels (based on their power-spectrum) were excluded. The ECG and EOG were used to automatically detect eye-blinks and heartbeats and to then remove them with signal space projectors. The data were then visually inspected for artefacts (muscle artefacts, head movements), with problematic time segments being excluded from further analysis. The cleaned MEG data were down-sampled to 1000 Hz to reduce the amount of data.

A 5-minute empty-room recording with the same sampling rate of 2400 Hz and an anti-aliasing filter of 800 Hz, but with no subject present in the magnetically shielded room, was obtained on each recording day. The goal is capturing the sensor and environmental noise statistics. Based on these recordings the noise covariance matrices were calculated for use in the source estimation process.

Forward modeling of neural magnetic fields was performed using the overlapping-sphere technique implemented in brainstorm (Huang *et al*., 1999). For the cortically-constrained weighted minimum norm estimate (wMNE) the lead-fields were computed from elementary current dipoles distributed perpendicularly to the individual cortical surface (Baillet *et al*., 2001). The individual surfaces were extracted with Freesurfer (version 5.3.0) using a tessellation of 15,000 (https://surfer.mnr.mgh.harvard.edu).

The MEG data preparation described in this section was common for the two compared approaches.

#### 2.1.1. Extracting the MEG resting state networks

Once the source-level data had been constructed, we extracted the MEG resting state networks. We used the megPAC approach as described in Florin and Baillet (2015) and the Envelope-ICA approach by Brookes and colleagues (2011). Both RSN extraction approaches first operate on the individual source-reconstructed MEG-data. To project the data from the individual to the standard anatomy for the cortical source model Freesurfer’s coregistered spheres were used as implemented in brainstorm (Fischl *et al*., 1999).

##### megPAC approach

For the megPAC approach we used the exact same parameters as described in the paper (Florin and Baillet, 2015). First, for each source time series of each subject the low-frequency phase that couples most strongly to the high gamma amplitude from 80-150 Hz was determined based on a phase-amplitude coupling measure (Ozkurt and Schnitzler 2011). Figure 1 shows the average low-frequency across subjects that exhibited the maximal phase-amplitude coupling to the gamma amplitude in each subject. Similar to previous results, the low-frequency was in the delta/theta range with no clear spatial pattern (Florin and Baillet, 2015). Both the phase and amplitude were extracted with a chirplet transform (Mann and Haykin, 1995), with a chirp factor of 0. For the low frequency corresponding to the identified phase the peaks and troughs were identified and the gamma amplitude (80-150Hz) was interpolated between these events. Through this process, for each source a new time series is obtained. These resulting time series were down-sampled to 10 Hz and then projected to the Colin27 brain. Within the Montreal Neurological Institute (MNI) space the cortical time-series from all subjects were first spatially smoothed (5mm Gaussian Kernel) and then concatenated. Subsequently, the spatial correlation matrix between all time-series was calculated. Finally, the resting state networks were determined as the principal spatial modes based on a singular value decomposition. These resting state networks were compared to the fMRI resting state networks obtained from the same subjects.

**Figure 1:**
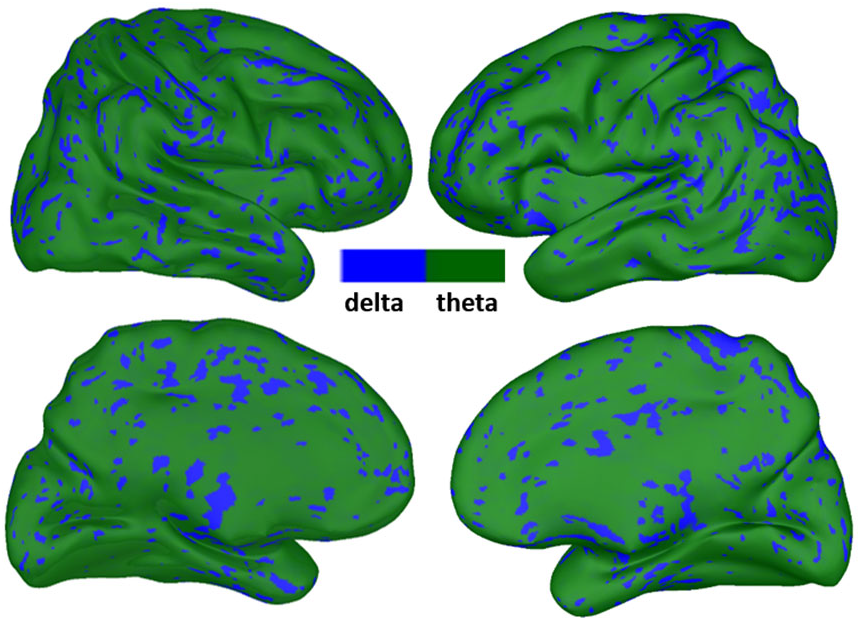
Low frequency of the megPAC approach. The cortical distribution of the low-frequencies averaged across subjects is displayed. This is the frequency that exhibited the strongest coupling to the high gamma amplitude and was subsequently used to extract the resting state networks with the megPAC approach.

##### Envelope-ICA approach

For the Envelope-ICA approach by Brookes and colleagues (2011) the Hilbert envelope is calculated in the 5 most common electrophysiological frequency bands: delta (1-4 Hz), theta (4-8Hz), alpha (8-12 Hz), beta (12-30 Hz), and gamma (30-50 Hz). We restricted our analysis to the approach by Brookes et al. (2011), except for also testing the effect of variations in filter settings and the ICA approach. To extract the envelope data for these five frequency bands we tested 2 different variants of filtering the data to obtain the Hilbert envelope:

1. Wide band: Filter width based on the pre-defined frequency bands (delta, theta, alpha, beta, and gamma). After filtering, the data were Hilbert transformed to obtain the envelope. This is the original approach described in Brookes *et al*. (2011).
2. Small band: From 1-4 Hz we band-pass filtered the data in 1 Hz bins, from 4-30 Hz in 2 Hz bins, and above 30 Hz in 5 Hz bins. The reason for this approach is that, based on the typical 1/f characteristics in wide frequency bands, the lower frequencies will dominate the envelope estimation. The resulting filtered time-series were Hilbert transformed to obtain the envelope. To obtain a single time series for the whole frequency band, we standardized the envelope data by z-scoring for each frequency bin and then averaged across the frequency bins within the respective band.

For each of the two binning approaches to obtain the Hilbert envelope we tested two different filter types: an infinite impulse response (IIR) filter and a finite impulse response (FIR) filter, both as implemented in the brainstorm function bst_bandpass_filtfilt. Both the IIR and the FIR filter were tested with the wide band and small band, resulting in four different filter combinations: small band FIR, small band IIR, wide band FIR, and wide band IIR filter.

Once the Hilbert envelope data at the individual level were obtained for each of the 4 filtering combinations, they were down-sampled to 1 Hz and projected onto the Colin27 brain. On the template brain the individual cortically-constrained data were spatially smoothed with a 5 mm Gaussian Kernel and then z-scored in the time-domain. The data from all subjects were subsequently concatenated in each frequency band. To extract the RSN, we follow Brookes et al. (2011) and employ the fastICA on the pre-whitened data based on a dimensionality reduction to 30 principal components. Because the reliability and numerical stability of one single ICA-run is usually not known, we performed an additional analysis using the ICASSO algorithm to extract the temporal independent components of each frequency band (Himberg *et al*., 2004). In the ICASSO algorithm the fastICA was run 50 times on the pre-whitened data. For the pre-whitening a principal component analysis was used to reduce the dimensionality to 30 components. The resulting independent components from each ICASSO run were then clustered based on the absolute value of the linear correlation coefficient between components. For these clusters the centrotype, which is the estimate that best represents all other estimates in the same cluster, was estimated and used for further calculations. To obtain spatial resting state networks the temporal independent components (the centrotype in case of ICASSO) were correlated with the envelope data. These correlation maps for each of the analyzed frequency bands were compared to the resting state maps based on the fMRI data.

### 2.2. MRI data acquisition and extraction of resting state networks

All magnetic resonance imaging data were obtained using a Siemens 3T PRISMA scanner using a 64-channel head coil. The high-resolution T1-weighted images were acquired by applying a 3D MPRAGE sequence (TR = 2300ms, TE = 2.32ms, ES = 7.2 ms, FA= 8°, FOV = 230mm x 230mm, isotropic pixel resolution of 0.9 x 0.9 x 0.9 mm, slice thickness of 0.9 mm, 192 slices). Resting fMRI data were recorded with echo-planar-imaging (EPI) acquisition, (TR= 776 ms, TE = 37,4 ms, flipangle = 55°, resolution 2.0 x 2.0 x 2.0 mm, slice thickness of 2.0 mm, 72 slices). The resting fMRI scan lasted 30 minutes. Subjects were also asked to rest with eyes open and to fixate on a paper cross to reduce eye movement.

T1-weighted images were automatically pre-processed with Freesurfer version 5.3.0 (recon-all) in order to extract the brain and cortex surface; brain extraction performed with Freesurfer yielded better results in the differentiation of cortical areas from the skull than FSL-BET. The resulting skull-stripped T1-weighted datasets were used as reference images in MELODIC 3.0 (https://fsl.fmrib.ox.ac.uk/fsl/fslwiki/MELODIC) after affine registration to standard MNI 2 mm space via FLIRT, a registration tool in FSL 5.0 (Jenkinson and Smith, 2001).

Data pre-processing of the fMRI 4D images was further carried out with FSL tools and the results were visually inspected. The following pre-processing was applied for each subject using MELODIC’s Pre-Stats feature: head motion correction via MCFLIRT(Jenkinson and Smith, 2001); removal of non-brain areas using BET (Smith, 2002), spatial smoothing with a Gaussian kernel of full width at half maximum (FWHM) 4 mm; grand-mean intensity normalisation of the entire 4D dataset by a single multiplicative factor; 100s high-pass temporal filtering.

Registration of each subject’s fMRI data to that subject’s high-resolution structural image was carried out by using 6 degrees of freedom registration with FLIRT (Jenkinson and Smith, 2001). Registration to the high-resolution structural MNI-152-2-mm standard space was achieved by using FLIRT affine registration

For the cortically-constrained analysis, we used the binarized cortical mask obtained from Freesurfer for the MEG case to restrict the fMRI ICA analysis to those cortical areas. We chose to variance-normalise time courses to make sure that mere differences in the voxel-wise standard deviations do not bias the PCA step and ICA cost function. Consistent with our MEG analysis, 20 ICA components were computed on data temporally concatenated across subjects.

After ICA decomposition, we chose the standard threshold of 0.5 for the IC maps following the recommendations in FSL Melodic. A threshold level of 0.5 in the case of alternative hypothesis testing means that a voxel ‘survives’ thresholding as soon as the probability of being in the ‘active’ class (as modelled by the Gamma densities) exceeds the probability of being in the ‘background’ noise class (see https://fsl.fmrib.ox.ac.uk/fsl/fslwiki/MELODIC).

In order to compare the fMRI resting state networks with MEG resting state networks we finally registered IC components, located in the MNI-152-2mm standard space, to the Colin27 brain.

### 2.3 Comparison of the fMRI and MEG resting state networks

In accordance with the previous literature, we consider the fMRI RSNs to be the benchmark and judge the success of the MEG RSN identification approaches based on their ability to reproduce these fMRI networks. We use two different procedures to measure the spatial correspondence of fMRI and MEG RSN.

Our first approach for the identification of the best-matching MEG maps for both the megPAC and Envelope-ICA approach as compared to the fMRI maps relies on the spatial correlation between the MEG and fMRI RSN. For each fMRI RSN we selected the best matching MEG component with the highest spatial correlation.

In our second approach, we instead rely on a binary measure by first thresholding the data. For this calculation, the MEG maps were thresholded at an absolute value of 0.3. The fMRI outputs were thresholded using the probability maps produced by FSL. A threshold value of 0.85 was used. We used D, which evaluates the spatial overlap (cortical sources) between one MEG map and one fMRI map (Mesmoudi *et al*., 2013):

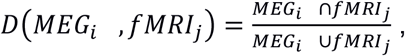

The overlap is calculated for each cortical source. If there is a perfect spatial overlap between a MEG map i and an fMRI map j, D will be 1. This is done for all MEG maps (i=1,…,20) and all fMRI maps (j=1,…, 20). For each fMRI-map the MEG-map with the highest D was selected.

### 2.4 Statistics for the comparison of the MEG approaches

To statistically compare the different approaches to extract the resting state networks we bootstrapped the envelope time series of each frequency 100 times across subjects for the Envelope-ICA approach and the megPAC time series 100 times for the megPAC approach. Afterwards the networks were extracted for each method as described above and the two metrics (spatial correlation and D) were computed for each of those repetitions. This provides a statistical distribution that captures sampling variability and allows for statistical inference. We first used a 2-way ANOVA to determine whether the bandwidth of the filter or the chosen filter type have a significant influence on the network estimation when employing the Envelope-ICA. Using post-hoc tests we then identified significant differences based on the filter choice. To compare the Envelope-ICA and the megPAC we used the best filter setting for the Envelope-ICA and performed a one-way ANOVA with a post-hoc test to determine the method with the highest spatial correlation / D for each network.

## 3. Results

In total, we extracted 20 ICs from the fMRI and MEG data. From the fMRI data we used 7 resting state networks for the further comparison: the frontal, parietal, left and right front-parietal, motor, visual, and default mode network. As an example, we show 4 of those networks along the rows of figure 2. The left column depicts the fMRI network, the right column the best corresponding MEG network according to spatial correlation. As can be seen from this figure, the MEG networks resemble the fMRI networks, but usually have a higher spatial spread than the fMRI networks. In the following, we will quantify in more detail the correspondence as well as several crucial choices for the Envelope-ICA implementation.

**Figure 2:**
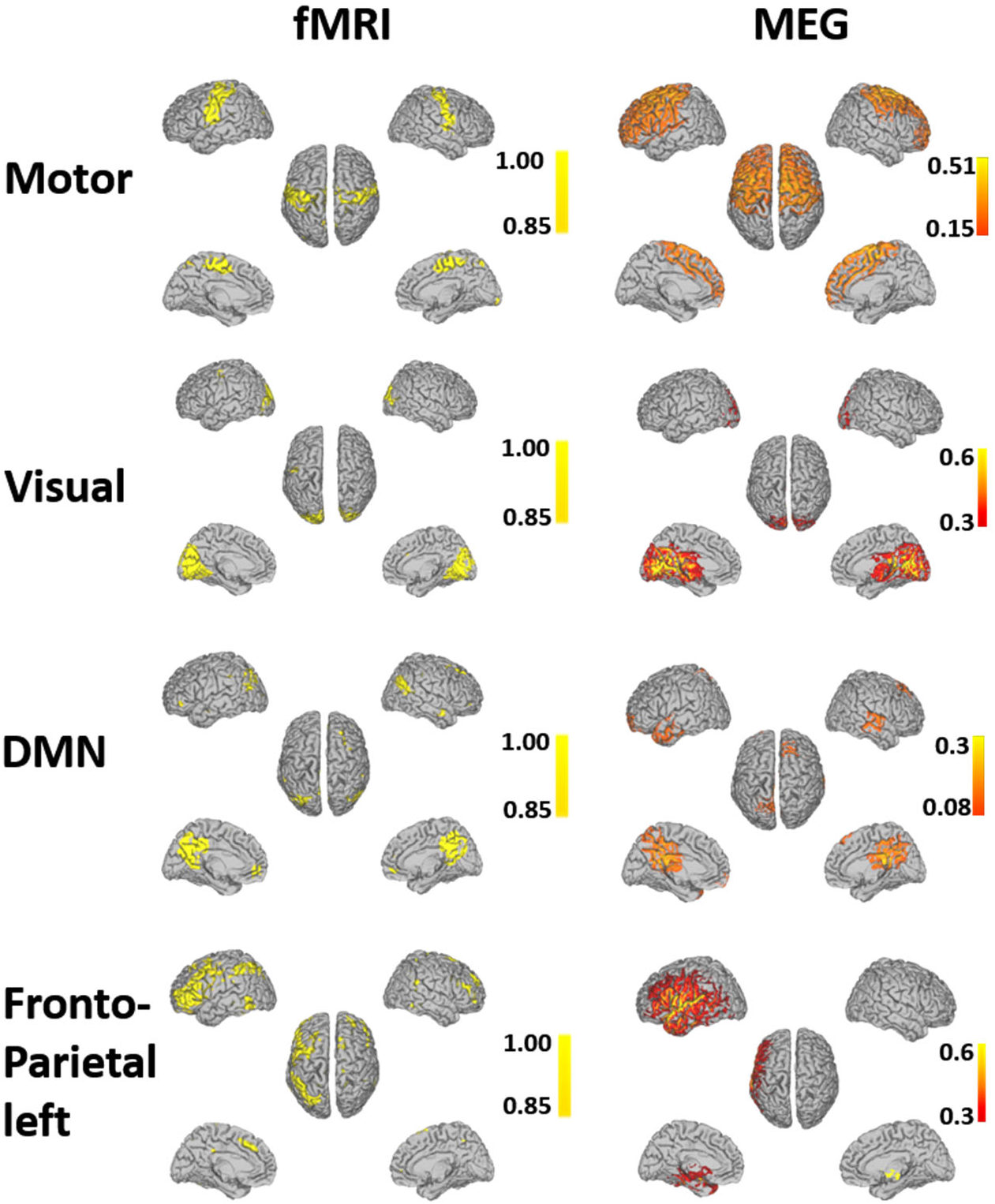
Highest correspondence resting state networks obtained from MEG and fMRI recordings. Note: The MEG resting state network with the best correspondence to the fMRI network according to the spatial correlation is shown. The motor network is based on the alpha envelope, DMN on the theta envelope, and the visual and fronto-parietal network on megPAC.

### 3.1 Method comparison for resting state network extraction

When extracting the Hilbert envelope for the Envelope-ICA approach different choices for the filter settings can be made. As described in the methods section, four different filter settings were tested. Using an ANOVA we identified that the filter type had a significant influence on the spatial correlation for the DMN, motor, visual, and frontal network (p<0.001). This was accompanied by a significant interaction with the chosen frequency band. Post-hoc analysis revealed that the narrow band filter yielded significantly better correspondence to the fMRI resting state maps for 1 network, while the wide band one yielded significantly better correspondence for 1 network (p<0.05). For the remaining networks 4 had a better, but not significantly better, correspondence with the wide band filter. Concerning the use of an IIR or FIR filter, the FIR filter led to significantly better results for 6 out of 7 networks, although the improved correspondence was only significant in one case (p<0.05). In the case of one network the IIR filter yielded significantly better results (p<0.05). Therefore, all following results will be presented for a wide band FIR filter.

When using the filter setting that uniformly performed best across frequencies (wide band FIR filter) for the Envelope-ICA approach the megPAC approach showed significantly higher correspondence in 4 out of the 7 cases to the fMRI networks (p<0.05). The Envelope-ICA approach had a significantly higher overlap for the parietal network in the beta band, the motor network in the alpha band, and the default mode network in the theta band (p<0.05). The spatial correlation values are plotted in figure 3.

**Figure 3:**
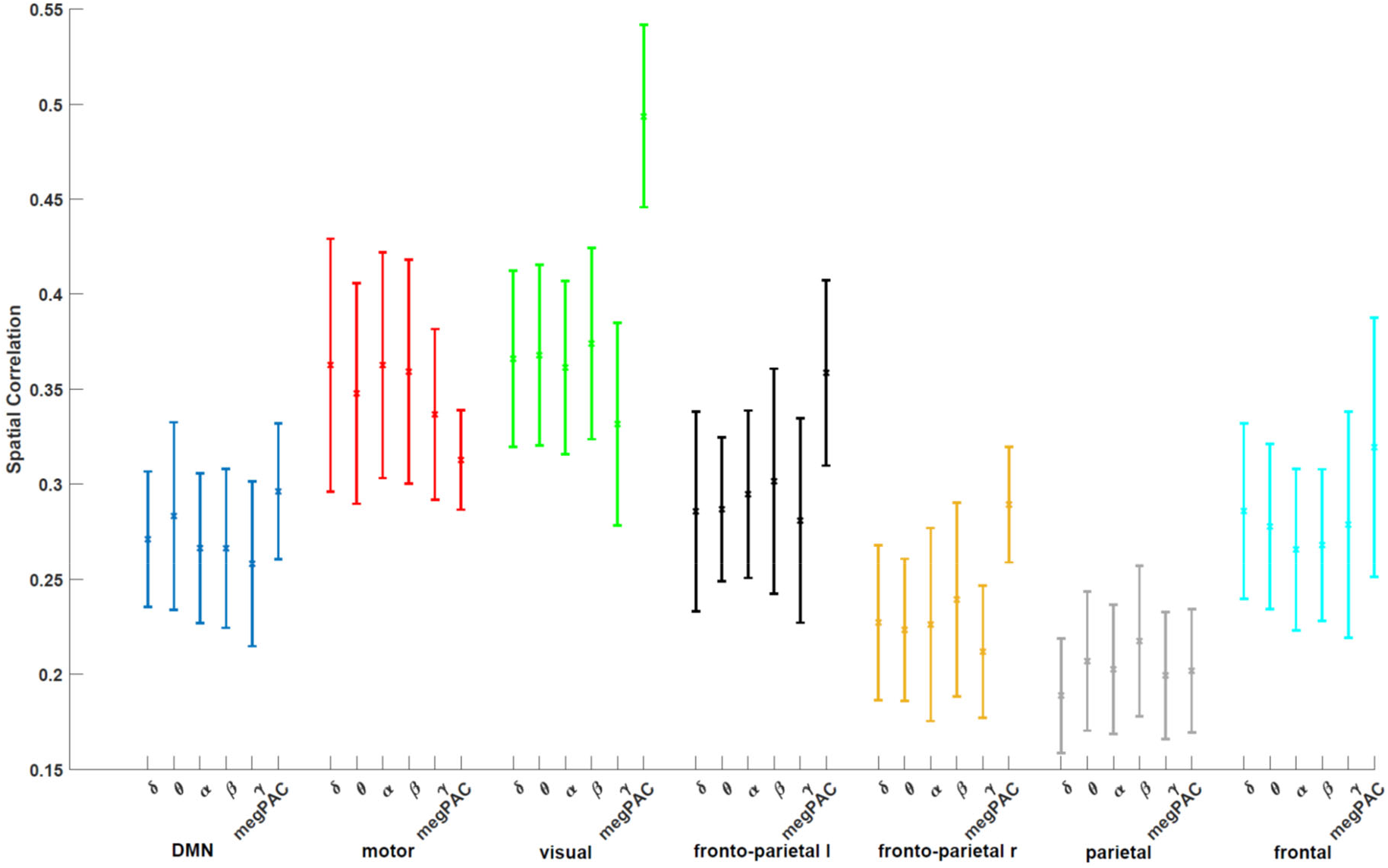
Spatial Correlation between fMRI and MEG RSN. The Envelope-ICA based results were obtained with the optimal filter setting of a wide band FIR filter. The different frequency bands for the Envelope-ICA approach as well as the megPAC approach (last value) are plotted on the x-axis. On the y-axis the spatial correlation is plotted between the fMRI resting state network and the corresponding MEG resting state network for the method provided on the x-axis. The error bars indicate the standard deviation.

In Figure 4 the D measure is used to compare the different MEG resting state approaches (for details see methods). There was a significant influence of filter type for all 7 networks (p<0.0001). Post-hoc tests revealed that the wide band filter showed significantly better correspondence between fMRI and MEG RSNs for 4 out of 7 networks and the FIR filter was better for 7 out of 7 networks (p<0.05). Comparing the RSN extraction methods using D, the megPAC performed best for all 7 networks (see Figure 4).

**Figure 4:**
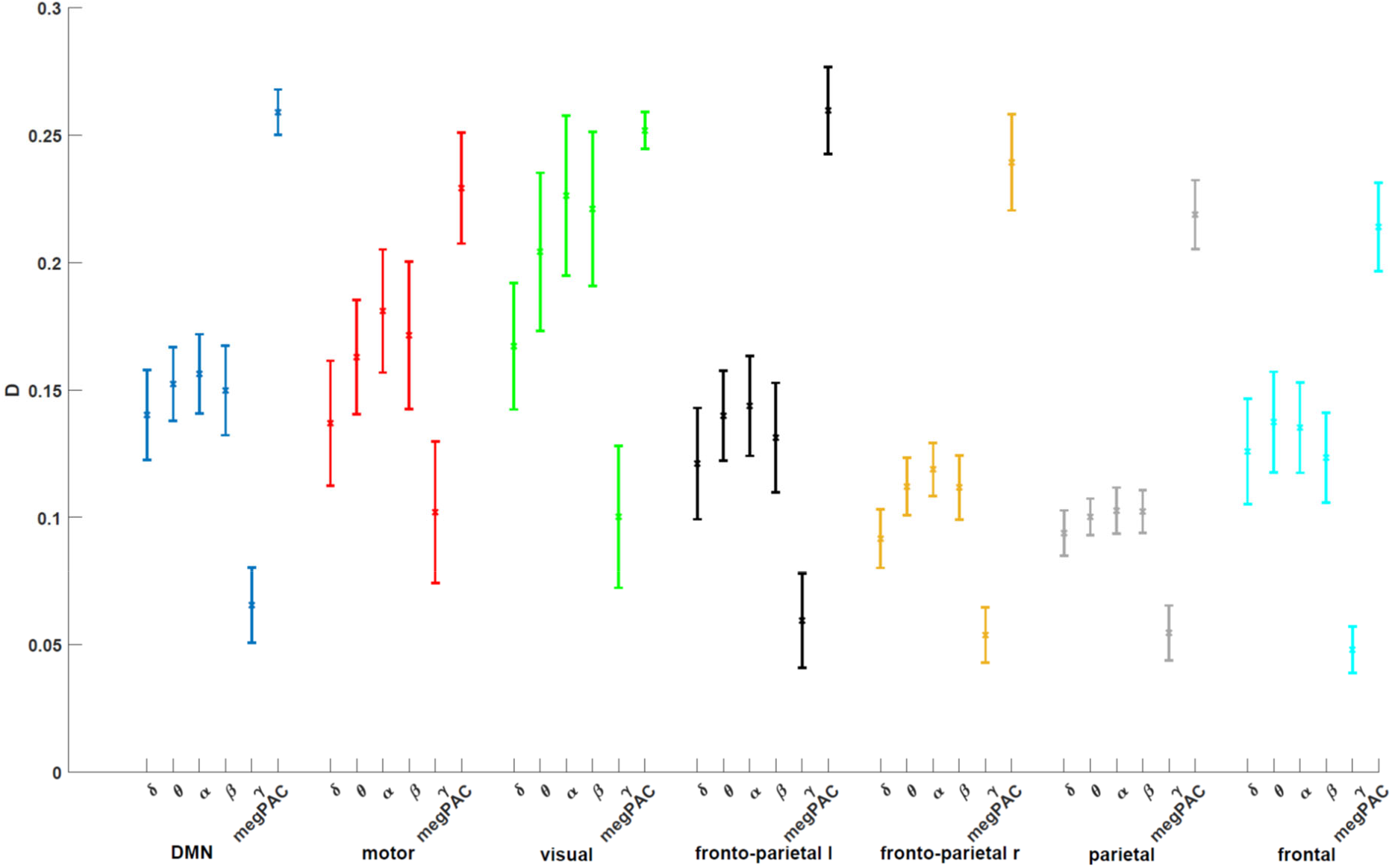
D between fMRI and MEG RSN. The Envelope-ICA based results were obtained with the optimal filter setting of a wide band FIR filter. The different frequency bands for the Envelope-ICA approach as well as the megPAC approach (last value) are plotted on the x-axis. On the y-axis D is plotted between the fMRI resting state network and the corresponding MEG resting state network for the method provided on the x-axis. The error bars indicate the standard deviation.

### 3.2 ICASSO for Envelope-ICA

The Envelope-ICA requires identifying Independent Components for each frequency band. However, as ICA is a high-dimensional optimization problem, the reliability of a single ICA run is not known (Eriksson and Koivunen, 2004; Hyvarinen, 2013). Therefore, to stabilize the results, we ran the ICASSO algorithm four times for each frequency band. Computationally, this is very expensive. One ICASSO run required two weeks of computation for each frequency band on a High-performance cluster with 200 GB RAM and 2 cores. Parallelization is complicated due to the large amount of RAM required. We then investigated whether the frequency components characterizing the individual networks are consistently the same across repetitions. When determining the IC map best matching the fMRI map, the spatial correlation value varied, resulting in ICs from different frequency bands showing the best correspondence to a particular fMRI resting state network. An example is provided in Figure 5 for 4 RSNs when using the wide band FIR filter settings. Within this figure, the spatial correlation between the fMRI RSNs and these 4 RSNs obtained with Envelope-ICA for the 5 frequency bands are plotted. For each RSN, the spatial correlation values for the four ICASSO repetitions are provided. As can be seen from the figure, these are highly variable across repetitions. Given these variable results, assigning the best match to a particular frequency band seems arbitrary.

**Figure 5:**
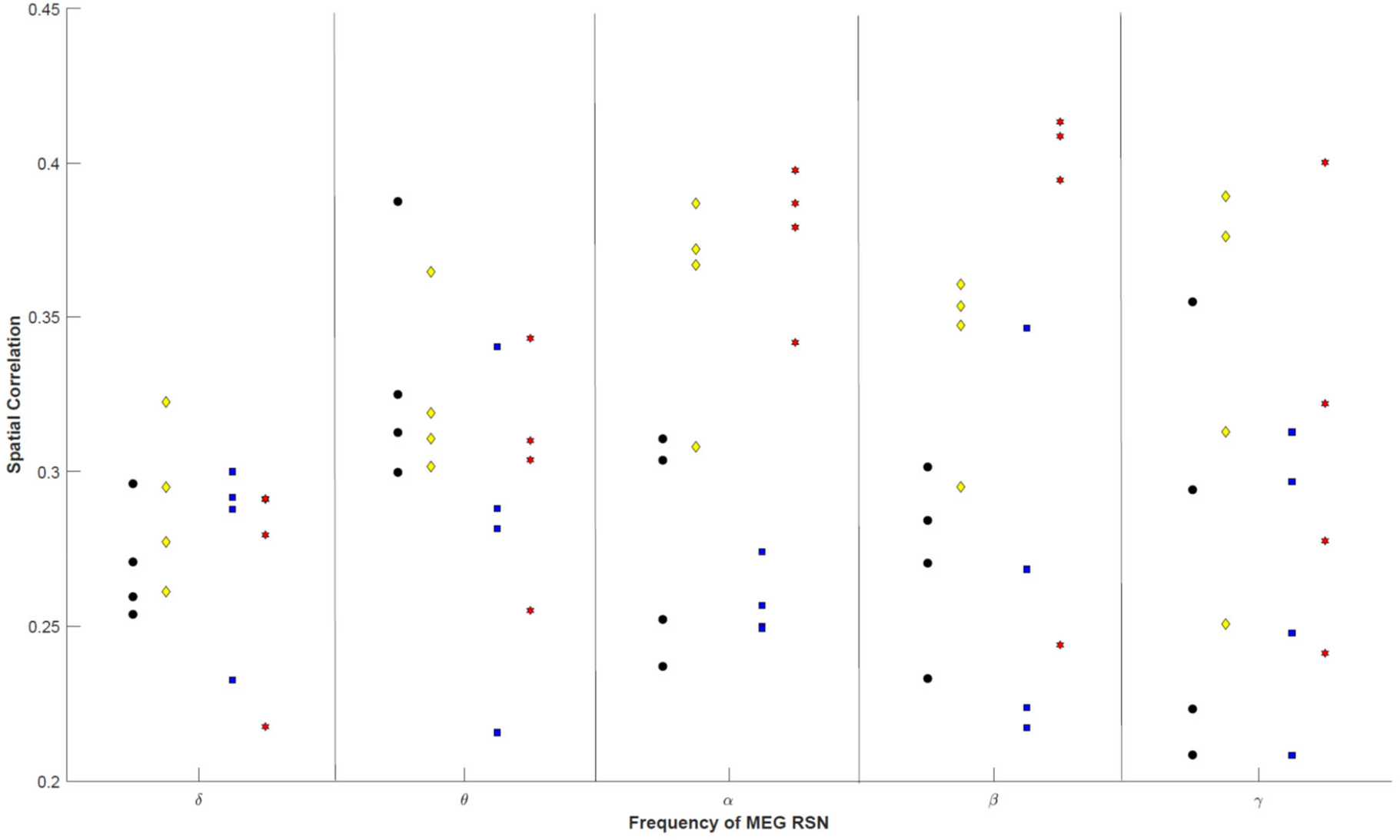
ICA-induced variability of spatial correlation for the Envelope-ICA approach for 4 resting state networks (colour coded). Within each frequency band, four repetitions of the ICASSO algorithm were conducted. The spatial correlation between the fMRI and MEG map is displayed. Note the variability across repetitions (black: DMN, yellow: motor, blue: fronto-parietal left, red: visual). The different frequency bands for the Envelope-ICA approach are plotted on the x-axis. On the y-axis the spatial correlation between the fMRI resting state network and the corresponding MEG resting state network is plotted.

## 4. Discussion

Within the present paper, we compared data-driven approaches to extract resting state networks as similar as possible to fMRI from MEG data. In order to minimize distortions introduced by RSN variability at the individual level, the comparison was made on recordings of the same subject once in the MEG and once in the fMRI. This is in contrast to previous studies employing standard fMRI maps for their comparison (Brookes *et al*., 2011; Florin and Baillet, 2015). The performance of MEG-RSN approaches was evaluated based on the spatial correspondence to the fMRI RSN maps. Concerning the use of a method the correspondence between fMRI RSN maps and MEG RSN maps was significantly better with the megPAC approach for 4 (7 in the case of D) of the 7 tested networks. Furthermore, when employing the Envelope-ICA approach the filter settings also affected the results: A FIR band-pass filter provided the best results. Concerning the physiological underpinnings of the fMRI RSN, we demonstrate that the frequency-specificity of the resting state networks proposed in the previous literature is not a given based on the Envelope-ICA approach.

### Frequency-specificity of the Envelope-ICA results

In the previous literature on Envelope-ICA-based resting state network extraction from EEG or MEG, the activity of each RSN network was ascribed to a particular frequency band within each study – with findings not always converging across studies. For example, the default mode network has been ascribed to both alpha and theta frequencies (de Pasquale *et al*., 2010; Brookes *et al*., 2011). When looking at the bootstrapped results in figure 3 and 4, it was not the case that there was a single dominant frequency band for each RSN considered. When we investigated this variability across repeated runs on the same subjects, such frequency-specific networks were not reproducible across different runs of the ICA algorithm with the same data. Thus, conflicting previous findings reporting different frequency components for one RSN may at least partially be explained by the variability in the ICA due to the dependence on the initial seed for optimization (Hyvarinen, 1999). Even using a stabilization procedure such as ICASSO (Himberg *et al*., 2004) did not improve the reproducibility across ICASSO runs. For ICASSO, our chosen number of resampling cycles of 50 might seem low. However, based on the recommendations for ICASSO and considering computational limitations of the current ICASSO implementation, 50 resampling cycles with 20 independent components produces 1000 estimates. It is quite unlikely that increasing the number of repetitions further would yield one dominant frequency band that did not appear in the 1000 repetitions before. This could also be an indication that each resting state contains information across a number of bands. This interpretation would also allow to reconcile previous heterogeneous results on frequency-specificity of the resting state networks (de Pasquale *et al*., 2010; Brookes *et al*., 2011; Hipp *et al*., 2012). Our results indicate that different frequency components can be identified as the best match for a RSN just because i) there was a different initial seed for the ICA and ii) the RSNs obtained from different frequency bands do not differ too much.

In addition, one has to keep in mind that the current ICA approach enforces independence in the temporal domain but not in the frequency domain. Therefore, this approach may be ill-suited to identify frequency-specific resting state networks. To obtain frequency-specific information one should aim for independence in the frequency domain and factorize appropriate matrices to yield spectro-spatial components (Hyvarinen *et al*., 2010). Future work should investigate in this conjecture.

### Choice of filter for Hilbert envelope

Before calculating the Hilbert transform for the Envelope-ICA, the data have to be band-pass filtered within the frequency bands of interest. When doing so the question arises whether the band-pass is chosen in small enough frequency bands to correct for the natural 1/f power decrease in MEG data. Within our tests, using a wide band filter provided better results. Moreover, the use of a FIR filter provided better results than an IIR filter in almost all cases, although the difference was only significant in one case.

### megPAC and comparison to previous studies

Recently, it was shown that phase and amplitude both contain information relevant for understanding the resting brain (Siems and Siegel, 2020). These findings could explain why the megPAC approach yielded better correspondence to the fMRI resting state networks than the Envelope-ICA approach. Using the Envelope-ICA approach it became apparent that one network cannot be easily ascribed to one dominant frequency band, suggesting that the other frequency bands may contain important information for their identification. Moreover, the envelope-ICA only relies on amplitude information. This might explain why the combination of phase and amplitude with the megPAC approach provided a better correspondence to the fMRI networks: the canonical resting state networks seem to be shaped by a low-frequency phase as well as the gamma amplitude. It is also worth pointing out that the Envelope-ICA approach only focuses on frequency bands up to beta.

A further advantage of the megPAC approach could be the computation time. Running ICASSO required two weeks of computation time on a high performance computing cluster for each frequency band. In contrast, running the RSN extraction with the megPAC approach only takes 1-2 hours on a standard computer. Therefore, from a computational and reproducibility point of view the megPAC approach seems more favourable.

There have been several studies that investigated the test-retest reliability of different estimates of resting state connectivity (Colclough *et al*., 2016; Garces *et al*., 2016; Dimitriadis *et al*., 2018). These studies have provided important guidelines on the choice of connectivity measures. Overall, amplitude-correlation based methods were more reliable than purely phase-based methods. Compared to our study a direct comparison to the fMRI resting state networks has been missing, i.e. the aim of those older studies was not to identify networks as similar as possible to the fMRI resting state networks but to provide guidelines on connectivity measures in general. This is also the reason why we had to limit our comparison to envelope-ICA and megPAC.

At the same time, the focus of our study was on data-driven methods as seed-based approaches necessarily involve a human element that is impossible to control for. Additional data-driven approaches involve finding time-resolved networks (Cribben *et al*., 2012; Baker *et al*., 2014; Yaesoubi *et al*., 2015; Vidaurre *et al*., 2018; Yaesoubi *et al*., 2018; Shappell *et al*., 2019). The methods used to study time-resolved activity vary widely in their underlying statistical assumptions as well as biological details (Lurie *et al*., 2020). Furthermore, source level network estimation using the time-resolved methods employs a limited number of regions of interest based on atlases. Both the spatial reduction and statistical assumptions of the time-resolved methods make them difficult to compare with ICA-based fMRI networks. Moreover, as the extraction of the canonical fMRI networks is not time-resolved, our aim here was to identify first markers of these resting state networks before investigating the time-resolution.

## Conclusion

In summary, it is possible to extract MEG resting state networks that correspond to the known canonical fMRI resting state networks. Of note here is that the spatial extent of the MEG resting state networks was larger than that of the fMRI networks. This was independent of the resting state approach used. Moreover, our results indicate one advantage of the megPAC approach compared to the Envelope-ICA approach. As the numerical estimation of ICA on real data introduces considerable variability, different ICA runs produced different dominant frequency-bands underlying RSN networks. This low frequency-specificity of the RSN when using the Envelope-ICA approach suggests that it may be problematic to ascribe activity within a given RSN to one particular frequency band. This finding may also hint at a way of reconciling conflicting findings in the literature on the main frequency components underlying individual RSN. This combination of different frequencies could also be the reason that the megPAC approach yielded the best correspondence for most networks.

## Acknowledgements

EF gratefully acknowledges support by the Volkswagen Foundation (Lichtenberg program 89387). Computational support and infrastructure was provided by the “Centre for Information and Media Technology” (ZIM) at the University of Düsseldorf (Germany). The authors would like to thank Johannes Pfeifer for his critical review of the manuscript and fruitful discussions.

